# Telomere-to-telomere genome assembly of Microsporidia sp. MB, a microsporidian symbiont of *Anopheles coluzzii* isolated from Burkina Faso

**DOI:** 10.64898/2026.02.24.707486

**Authors:** Roland Pevsner, Julien Martinez, Deepak-Kumar Purusothaman, Beth C. Poulton, Abdelhakeem I. Adam, Ewan R.S. Parry, Saré Issiaka, Abdoulaye Diabaté, Steven P. Sinkins

## Abstract

**Background:** Microsporidia sp. MB is an intracellular parasite of anopheline mosquitoes identified across the African continent. Microsporidia sp. MB infections appear to have no significant effect on host fitness and vertically transmit, whilst infected individuals have exhibited significantly reduced levels of *Plasmodium falciparum* transmission. These combined characteristics make Microsporidia sp. MB a promising candidate for use in malaria control. A comprehensive genome would greatly facilitate investigation into the evolution and biological pathways underlying important phenotypes, such as the mechanism of malarial inhibition.

**Results:** In this study, we present the *de novo* assembly of the first complete genome of Microsporidia sp. MB SOUVK7. Multi-platform sequencing was performed on ovary samples of laboratory established *Anopheles coluzzii* collected from Burkina Faso. The SOUVK7 genome has a total size of 9.16Mb, encodes 2,435 genes and is organised into 13 chromosomes with telomeres identified at all flanks. Telomeric repeats exhibit a 4-mer motif due to a glutamine deletion previously unobserved in Microsporidia. Ploidy analysis of Illumina reads predicts MB as tetraploid, whilst analysis of CpG methylation and retroelements highlights loci in all chromosomes with characteristics consistent with regional centromeres. Orthology analysis identifies several key genes in pathways associated to telomeric and centromeric maintenance, along with methylation and host invasion machinery. A loss of several components of the infection machinery is observed in Microsporidia sp. MB and the wider Enterocytozoonida, consistent with a general trend towards genome size reduction in the clade.

**Conclusion:** This study provides the first complete, telomere to telomere assembly of Microsporidia sp. MB, offering new insight into the genomic architecture of Microsporidia sp. MB and the broader Mrazekiidae family.

## Background

Pathogenic species of the Microsporidia are emerging concerns in public health, sericulture and aquaculture sectors [1–3]. This phylum of obligate intracellular parasites infect a broad range of hosts across all animals, exhibiting a variety of symptoms from digestive distress to growth retardation and high mortality. Due to the opportunistic pathology of microsporidia, very few symbiotic relationships have been reported with their hosts. Microsporidia sp. MB (MB) was recently discovered in several species of *Anopheles* mosquitoes in Kenya and has also been reported in wild populations across sub-Saharian Africa [4–7]. In Kenya, mosquitoes infected with MB and challenged with malaria-infected patient blood had reduced rates of *Plasmodium* infection in their midgut and salivary glands when compared to MB-negative mosquitoes. This data wais corroborated by a lack of MB/*Plasmodium* coinfections reported through extensive field sampling of wild *Anopheles* mosquitoes. These results strongly indicate a malarial inhibition effect with *Anopheles* mosquitoes, provided by MB. As MB appears to confer no significant costs to host longevity or fecundity, transmits maternally and exhibits a symbiotic or commensal relationship, MB shows great potential as an alternative agent in malaria control.

Microsporidia exhibit extensive levels of genomic reduction of both coding and non-coding regions, believed to be the result of their obligate parasitic lifestyles [8]. For this reason, Microsporidia species still have the smallest reported complete genomes of any eukaryotic organism. The reduction in total genome size is also reflected in low total gene counts of approximately 2,000 to 6,000. Phylum-wide analysis has previously suggested that approximately 48% of gene families within Microsporidia are unique to their phylum and are often even specific to families or clades. As observed in other intracellular parasites, Microsporidia genomes are also distinguished by significantly lower GC contents of 19 to 55%, often making them easy to differentiate from eukaryotic host reads. Currently our understanding of microsporidian chromosomal architecture is limited due to the low number of published complete or chromosome level assemblies (NCBI: 1 and 6 assemblies respectively; as of July 2024). As research initiatives continue to assess MB’s potential as an agent for malaria control, a comprehensive and complete genome is needed to support investigation into the mechanisms of *Plasmodium* inhibition and host interaction, along with population genetics and diversity across tropical Africa. Four MB genome assemblies, primarily from isolate MB AHL03, have previously been released but are highly fragmented (≤2,400 contigs) and thus incompatible for analysis of genome synteny or chromosomal architecture [9]. These genomes are all assembled from separate collections in northern Kenya, and each likely represent populations with low genetic differentiation to one another, complicating efforts to identify effective genetic markers for population analysis.

In this article, we present the first complete *de novo* assembly of the Microsporidia sp. MB genome, which was isolated from laboratory established *Anopheles coluzzii* collected in Burkina Faso. This expands the diversity of available MB genomes but also acts as a new reference for investigation of microsporidian genome structure and architecture, and can aid reassembly of prior MB genome releases. Furthermore, given the low number of chromosome level and complete genomes within the Microsporidia phylum, this novel assembly expands our genomic understanding of the wider Enterocytozoonida clade [10].

## Results

PacBio sequencing of *An. coluzzii* ovaries infected with MB generated 766,153 reads with an average read length of 15.5 kb (max length: 44.5 kb). Total GC content showed two distinct read populations of approximately 31% and 45%, hypothesised as MB and host, respectively. Upon removal of host reads, 125,854 reads remained (16.4% of total reads) with a GC content of only ∼31%, suggesting the 45% GC reads belonged to the host (**S*1*** Fig: A & B). This reduction in reads had little effect on mean read length but led to a notable drop in maximum read length (15.3 kb mean length; 29.8 Kb max length). Illumina sequencing provided a dataset of 105,381,324 paired reads with an average sequence length of 93.8 bases. After removal of host and contaminant reads, 13 million reads remained and accounted for 12.4% of total reads.

A preliminary assembly of non-host PacBio reads using Hifiasm in default diploid mode yielded 77 contigs, 56 of which were assigned to Microsporidia by taxonomic profiling (S1 Fig). Plotting of this assembly showed 13 clusters of duplicated contigs, strongly suggesting that the contigs were incorrectly phased and likely due to genome polyploidy (**Fig 1**.A). GenomeScope analysis of both Illumina and PacBio reads predict a tetraploid kmer distribution with significant bias towards AAAB kmers over AABB kmers, and predicted haploid genome sizes of 8.3 and 7.2 Mb, respectively. SmudgePlot outputs also support tetraploidy with a majority prediction of AAAB kmers in Illumina reads (**Fig 1**. D).

**Fig 1:**
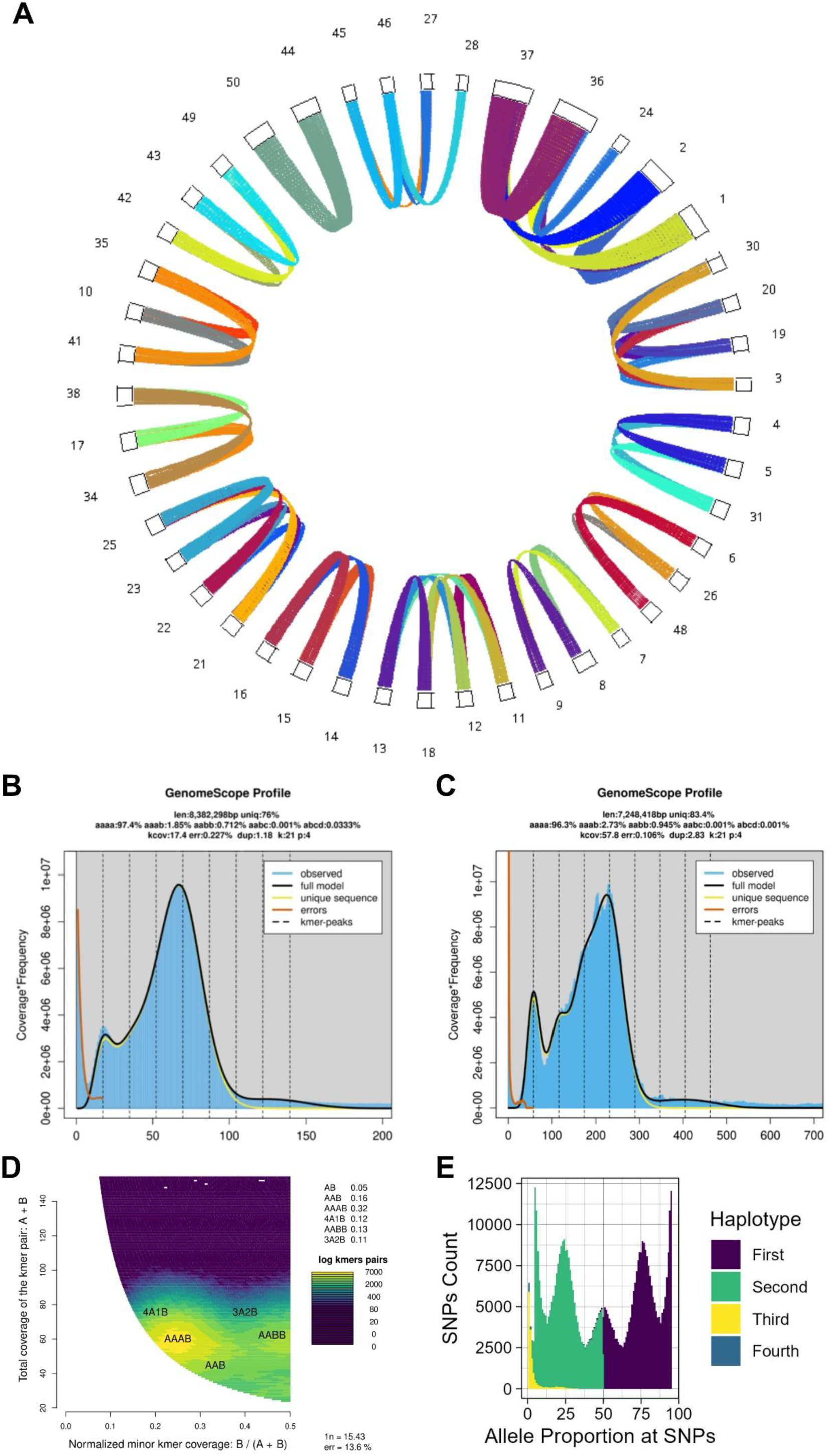
Predicting tetraploidy in Microsporidia sp. MB SOUVK7 through multiple analysis tools. (A) Chromosomal synteny of initial haploid assembly of MB SOUVK7, re-organised *to* highlight duplicate syntenic contigs. 13 distinct groups of syntenic contigs are identified with 2 to 5 contigs in all groups. The most abundant counts are 3 and 4 contigs encompassing 7 and 4 contig groups, respectively. Synteny analysis and visualisation was generated with MCScan. (B-C) GenomeScope plots of Illumina and PacBio reads, respectively. Although both plots only exhibit 2 clear peaks, additional 2 kmer peaks are predicted between the observed peaks. Both profile statistics also confirm tetraploidy with 4-mer haplotype predictions, additionally suggesting autotetraploidy with both profiles having higher AAAB heterozygous abundance. (D) SmudgePlot heatmap visualisation of PacBio read data highlighting the significantly elevated abundance of 4-mer pairs (AAAB & AABB) compared to other kmer values within the data. (E) Histogram of ploidyNGS results from PacBio read data, plotting distribution of reads associated to SNP haplotypes across the genome. Colour shows the order of haplotype abundance at SNPs. The five peak distribution of the plot at 5%,25%,50%, 75% & 95% clearly correlated to exemplary tetraploid results provided with the tool [11].

Reassembly of non-host reads assuming tetraploidy generated 20 contigs, with SprayNPray separating them into two distinct populations; a dense clustering of 14 contigs (GC content: ∼31%, read depth: ∼175-225x) and a sparser cluster of 6 contigs (GC content: 39-56.5%, read depth: ∼3-13x) of microsporidian and bacterial origins, respectively. BLAST searches of the bacterial contigs again found strong homology to C*edecea neteri*. Of the 14 remaining microsporidian contigs, one contig was identified as an outlier due to its lack of telomeres, significantly reduced size and elevated read depth (9.4kb and ∼245x, respectively).

The finalised genome had a total size of 9,162,986 bp (9.16Mb) across 13 complete chromosomes with an N50 of 650,044 bp (**Fig 2**). This was broadly consistent with the genome sizes predicted by GenomeScope during the ploidy analysis. Based on this final assembly, PloidyNGS plots also supported a tetraploid genome, with five clear peaks across individual and consensus plots of all chromosomes, consistent with example tetraploid data (https://github.com/diriano/ploidyNGS/blob/master/test_data/ploidyNGS_results/DataTestPloidy4.tab.ExplorePloidy.png). Chromosome length ranged from 477 kb to 1.6 Mb with a mean coverage of 208x and mean GC content of 31.13%. Illumina reads mapped against the genome with a mean depth of 172.7x. Funannotate and Barrnap predicted a total of 2,435 genes distributed mostly evenly across all chromosomes, comprising 2,294 protein-encoding genes, 87 tRNAs and 54 rRNAs. BUSCO analysis reported similar assembly completeness to other Enterocytozoonidae species with 98.4% of complete genes and 1.4% missing genes (**Fig 3**). The multi-gene phylogeny provided by OrthoFinder also placed Microsporidia MB SOUVK7 within the Enterocytozoonida, in immediate relationship to Microsporidia MB AHL03 and *Vittaforma corneae* (**Fig 3**).

**Fig 2:**
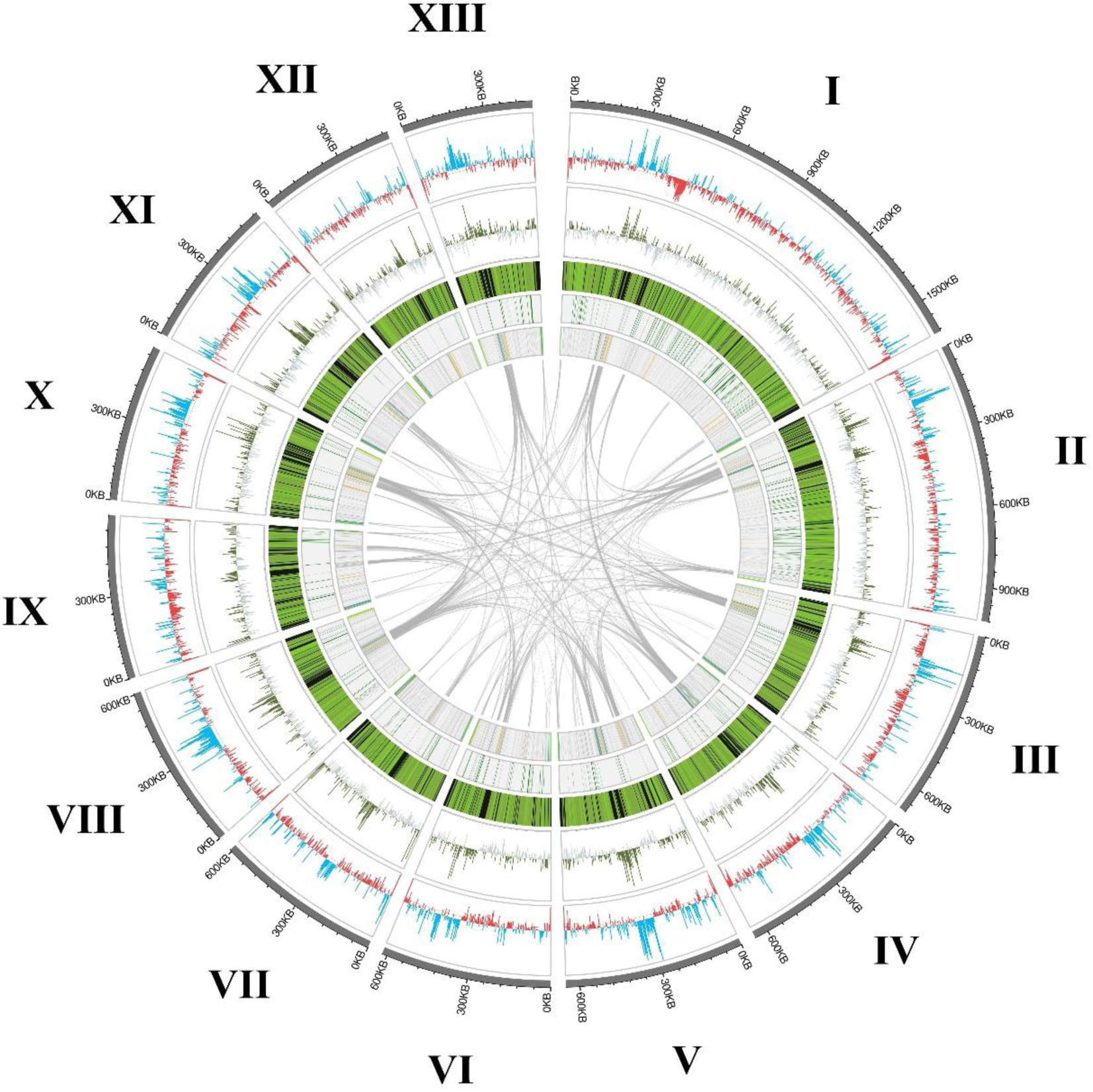
Circos plot of all chromosomes of Microsporidia sp. MB SOUVK7, highlighting genetic characteristics. The lengths of the 13 chromosomes are represented by their respective segment, with circles representing different analytical datasets. Outer circle shows local GC content across chromosomes as the mean across a 3,000 bp window. Cyan and red peak colours illustrate the windows GC content being above or below the mean GC content of the assembly (31.13% GC), respectively. The second circle plots peaks of methylation density as total CpG bases per 3,000 bp window. Green peaks represent windows of CpG density above the assembly mean (0.01306 CpG/base), whilst grey represents windows below mean. The third circle demonstrates the local gene density within the chromosome, calculated as the number of gene annotations per 3,000 bp window. Density is shown as a heatmap with black denoting zero genes per window, with continuous green gradient for windows with greater than one gene per window. The fourth circle represents the location of BUSCO genes across the genome with green, blue and red points representing complete, duplicated and fragmented genes respectively. The inner circle highlights the location of various retroelements, coloured by retroelement groups determined by RepeatModeler (cyan: LINEs [L1, RTE-X, RTE-BovB], orange: LTR [Gypsy], light grey: LINE [Dong-R4]; on a dark grey background).

**Fig 3:**
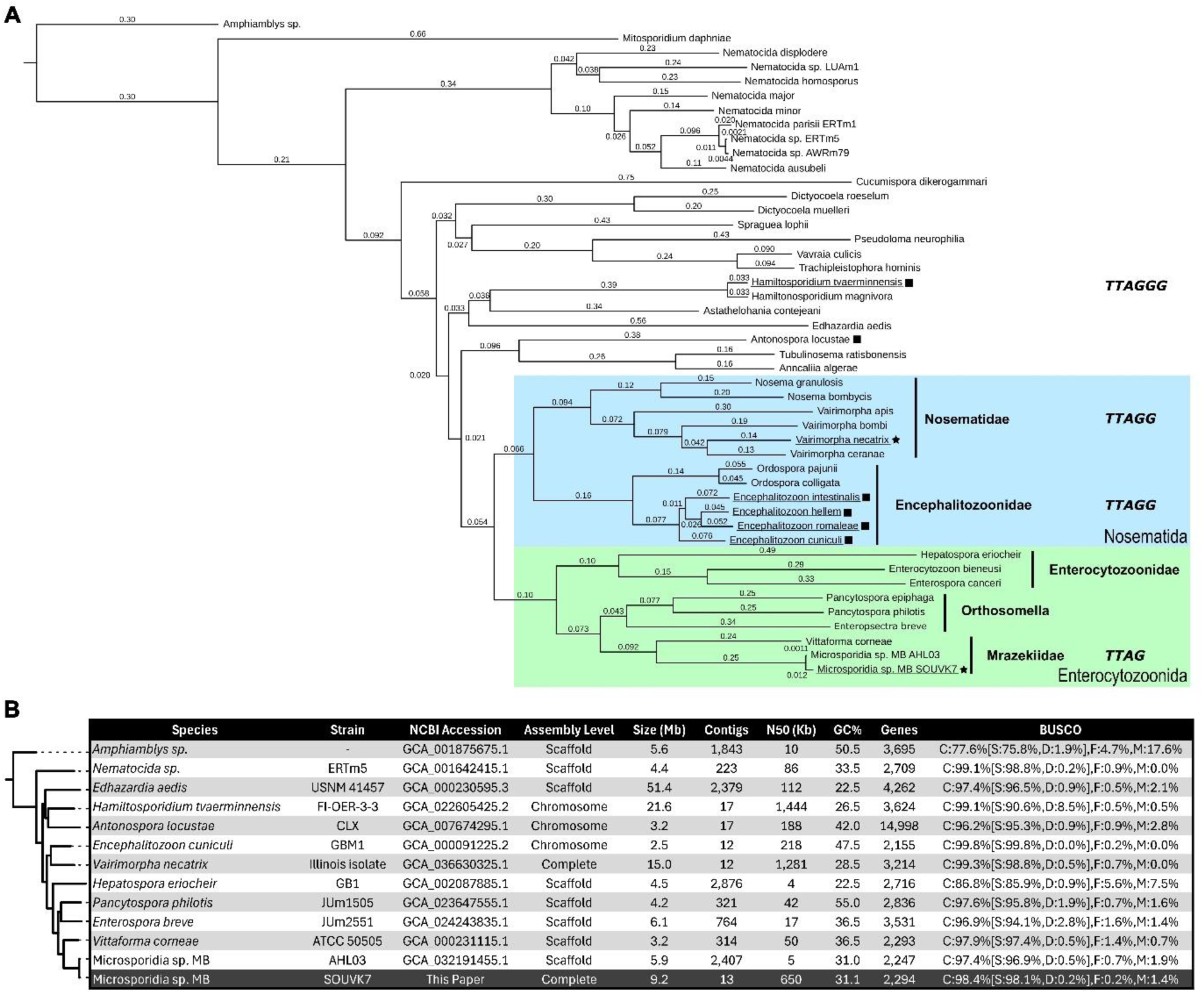
Comparative phylogeny and assembly statistics of MB SOUVK7 to reference genomes of the Microsporidia phylum. (A) Multi-gene phylogeny of the Microsporidia sp. MB SOUVK7 in the context of the Microsporidia phylum. Coloured boxes and vertical lines highlight clades and families of related species, respectively, as per Bojko et al. 2022. Solid square and star symbols indicate species possessing chromosome and complete level assemblies, respectively. Underlined species assemblies include telomeric kmer sequences, with encoded kmers in bold italics to the right of the respective species. Phylogeny was generated as part of the OrthoFinder pipeline and the tree was visualised with TreeViewer. Microsporidia sp. MB SOUVK7 localises closely the MB AHL03 within the Mrazekiidae, a clade within the Enterocytozoonidae family. (B) Comparison of the assembly characteristics for Microsporidia sp. MB SOUVK7 against other pertinent microsporidia species. Tree was generated from a multi-gene phylogeny generated by OrthoFinder, with Amphiamblys sp. as the outgroup and branch numbers representing length. Table values are collected from NCBI with BUSCO scores being calculated for total nucleotide sequence with the ‘microsporidia_odb12’ dataset.

During the masking step of assembly, RepeatMasker identified 40.66% of the genome as repeat sequences, predominantly as retroelements and unclassified repeats accounting for 26.17% and 6.73% of the total genome, respectively. The most abundant of these retroelements were Dong-R4 type long interspersed nuclear elements (LINEs) which evenly distributed within and between chromosomes (**Fig 2**). The remaining LINEs and long terminal repeats (LTRs) largely localised at the termini or discrete regions in the centre of chromosomes. Terminal regions of all MB chromosomes were found to possess simple 4-mer telomeric motifs of CTAA/TTAG. Telomere repeat regions varied in size between 1312bp and 7145bp, with a mean length of 3,900 bp and were often flanked by further LINE repeat groups. Telomeric regions exhibited significantly lower frequencies of methylated cysteines when compared to mean CpG methylation of the whole assembly (0.00326 vs 0.01306 CpG/base, respectively). Repeat-rich areas within the chromosomes, comprised mostly of L1 type LINEs and Ty3 type LTRs, spanned large central sections (30-82kb) of most chromosomes. Synteny and BLAST analyses of these repeat-rich regions showed low synteny conservation between chromosomes, however all regions consisted of conserved repeat motifs uniquely organised into each region. These regions were consistently the areas of highest local GC content within chromosomes (47.6% Mean local GC, 2.188% StDev). The high density of repeats was also reflected by decreases in local gene density. Although methylation was observed across the entire genome, these regions correlated with high CpG methylation. Most chromosomes possessed single examples of these select regions, however chromosomes II and VI potentially displayed pairs or splits in these regions. Gene annotation of SOUVK7 predicted 2,294 protein-encoding genes, and was comparable to other members of the Enterocytozoonida, but remained low for the broader phylum (**Fig 3**).

Different signal prediction tools identified between 133 and 462 genes with signal peptides and a total consensus of 12 genes, with DeepLoc and TargetP providing largest consensus of 160 genes (S4 Table). Similarly, comparison of transmembrane region (TMR) prediction tools showed a closer consensus to one another of 424 genes (18.7% genes; 521 & 430 predictions for DeepLoc and DeepTMHMM, respectively). Over 80% of gene products were predicted by DeepLoc as intracellular, whilst only 151 and 43 gene products are predicted to be extracellular or transmembrane (6.6% and 1.9%, respectively). 2144 of MB SOUVK7 genes (93.5%) are assigned to microsporidian orthologies with MB SOUVK7 appearing in 16.7% of all orthogroups and containing only 2 species-specific orthogroups. Furthermore, 1998 MB SOUVK7 genes were found to have at least one ortholog in MB AHL03, accounting for 95.6% of orthogroup-allocated genes. Based on these parameters and total gene counts, gene predictions of MB SOUVK7 and MB AHL03 were broadly considered homologous, as reflected in the resulting multi-gene phylogeny (**Fig 3**).

Homology searches for telomeric and heterochromatin maintenance components (GO:0000723 & GO:0034508, respectively) identified 24 MB orthologs of the 38 reported genes from *E. cuniculi*. Within the telomere maintenance machinery, only telomerase reverse transcriptase and the Rad32-Rad50 protein complex was found in MB, accounting for only 25% of *E. cuniculi* components. Notably, MB lacked components of both the Cdc13-Stn1-Ten1 and shelterin complexes, associated with protecting against telomeric degradation despite well-conserved motifs still being seen in all chromosomes. OrthoFinder searches for broader fungal GO components identified two additional orthologs of telomeric maintenance proteins in MB (RAD51 & RAD52)[12,13]. Although no components of ‘Centromere Complex Assembly’ (GO:0034508) were previously reported in Microsporidia [14], OrthoFinder found four associated orthogroups including hits for all species, including MB. Orthogroups were found to contain Ino80 and YTA7, MIS16 and MAM1 genes associated with centromere remodelling, histone chaperoning and meiotic regulation, respectively.

MB orthologs for ‘Heterochromatin Formation’ (GO:0034508) demonstrated the largest divergence from previous *E. cuniculi* predictions. BLAST searches only found 21 MB ortholog genes to the 30 *E.* cuniculi genes assessed. Most notably, MB only possessed three of five components of the CLRC complex required for heterochromatin formation (Swi6, Cul4 and Rik1), of which all five components were reported to be essential for CLRC function [15]. MB encoded orthologs for the four core histone proteins (H2A, H2B, H3 & H4) and significantly CENP-A, a highly divergent variant of H3 histones most commonly associated with epigenetic mechanisms in the centromere [16]. Finally, all methylation components in the Encephalitozoonidae were found in MB including all 3 paralogs of rRNA m5c methyltransferase, with the exception of NEP1 (rRNA SSU methyltransferase).

Spore wall proteins (SWPs) and polar tube proteins (PTPs) play essential roles in interaction and infection of host tissues. MB SOUVK7 possessed several proteins associated with the spore wall and polar tube infectious machinery. OrthoFinder results provided at least one Microsporidia orthogroup for 19 of the 24 SWPs and PTPs examined (S4 Table), with the remaining 5 search proteins being unassigned or only existing in single species orthogroups. An additional 7 protein groups (PTP7, SWP1, NbSWP5, SWP8, SWP25, SWP26 & SWP30) lacked species diversity outside of the Nosemida so were removed from the final results. The remaining protein families included 6 of each of the PTPs and SWPs with extensive coverage across the phylum (**Fig 4**.A). No orthologs were identified in the outgroup Metchnikovellid species of *M. daphniae* and *Amphiamblys sp.*, with the exception of aquaporin (NbAQP) which was found in all species except *V. apis* and *V. bombi*. The extreme fragmentation of these two genomes may have contributed to this lack particularly when compared to other genomes within the genus. Furthermore, *Nematocida* species possessed orthologs for only PTP3, EnP1 and SWP9, with further tBLASTn searches verifying the lack of all other gene families. Due to the basal nature of the Nematocida, this lack may have highlighted the limitations of OrthoFinder to identify orthologs within highly divergent species. This was shown as additional BLASTp searches were required to initially identify orthologs for EnP1 and SWP9, however the lack of detection of other gene orthologs may have been due to a lack of resolution in the phylogeny at the base of the phylum.

**Fig 4:**
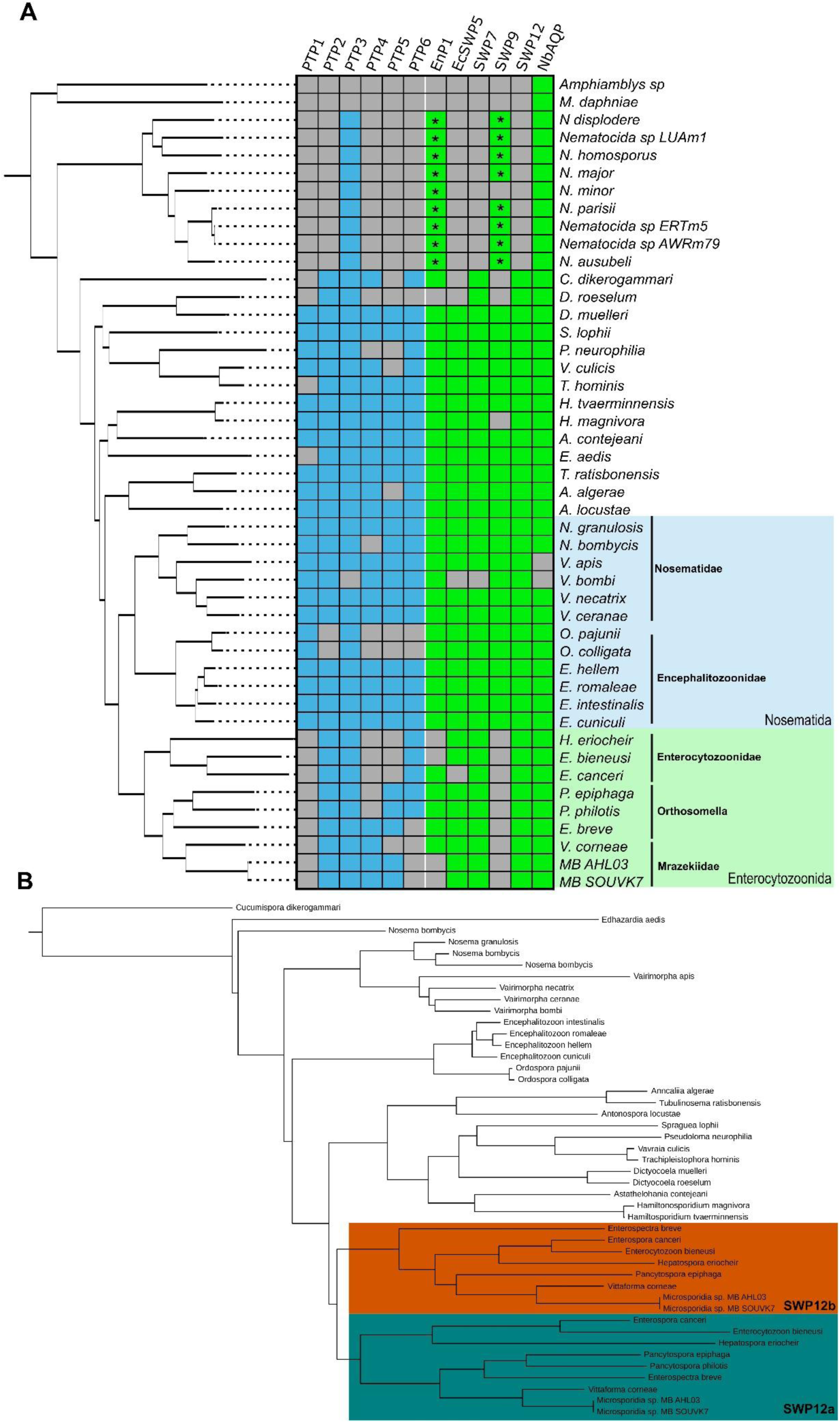
(A) Orthology of principal polar tube and spore wall proteins across the Microsporidia phylum. Phylogeny of all analysed species are illustrated to the left of the table and is based on the total species phylogeny generated by OrthoFinder. Species in possession of at least one ortholog of the associated polar tube and spore wall proteins are represented by blue and green colours, respectively. Species lacking any orthologs are illustrated by grey. Orthologs denoted with a star represent *Nematocida* orthologs first identified through BLASTp searches and verified to exist in a single orthogroup. **(B) Phylogeny of orthogroup associated with spore wall protein 12**. General phylogeny of the group follows the multi-gene phylogeny of the phylum (Fig 3), with a branch duplication at the Enterocytozoonida node. The two distinct isoforms are highlighted in green and gold for SWP12a and SWP12b, respectively.

Within the Enterocytozoonida, total loss of PTP1, PTP4 and SWP9 orthologs was observed across all 9 species including MB SOUVK7, despite near complete representation within all other species of the phylum. PTP4 and PTP5 families were both allocated to the same orthogroup, with one MB ortholog (ACK3DK_000951) allocated to the PTP5 family, due to its taxonomic position. As no orthogroups associated with PTP4 and PTP5 included any predictions for *V. corneae,* extra orthogroups for previously determined orthologs (VICG_01195 and VICG_01807, respectively) were further investigated [17]. The two orthogroups of PTP4 and PTP5 only included *V. corneae* orthologs and complimentary MB orthologs, suggesting a potential misidentification of the PTP orthologs in prior literature or prediction of an alternative PTP family uniquely associated to the Mrazekiidae.

OrthoFinder also identified orthologs of 4 SWP families (EcSWP5, SWP7, SWP12 and AQP) in both MB genomes, whilst reporting loss of SWP9 and EnP1. All SWP9 orthologs correlated to a single orthogroup and included all remaining species except all Enterocytozoonida species. Further tBLASTn searches confirmed these losses with no significant hits, supporting the family wide loss of this gene. No EnP1 orthologs were identified in either MB genome despite the detection of orthologs for all other Mrazekiidae species and this was also confirmed with no significant tBLASTn hits to either genome. The lack of EnP1 was not unique to MB however as orthologs were also missing in *E. bieneusi*, *H. eriocheir*, *D. roeselum* and the entirety of the *Nematocida* genus.

Searches for SWP12 presented two MB distinct paralogs within the same orthogroup whilst only single orthologs were identified across the phylum. This orthogroup also included both SWP12 paralogs for all Enterocytozoonidae species, but tBLASTn searches still failed to identify secondary paralogs across the rest of the phylum (**Fig 4**.B). The resulting orthogroup tree followed the standard Microsporidia phylogeny and showed a clear duplication event, resulting in the two Enterocytozoonidae paralogs, occurring at the basal node of the family [18]. Both branches mimicked that of the expected family phylogeny except for a single SWP12a ortholog in *P. philotis*. The comparison of taxonomic lengths between both branches suggested a closer conservation of the complete branch to the wider protein family, leading to the tentative naming of SWP12a and SWP12b to the complete and partial branches, respectively. Alignment of the paralogs showed strong conservation along the lengths of both genes, however SWP12a appeared to possess an additional 8-15aa at the N-terminus, likely contributing to much of the phylogenetic separation of the paralogs.

## Discussion

Despite the growing number of Microsporidia reference genomes on NCBI, very few are resolved to a chromosome or complete level. Furthermore, the taxonomic variety of these genomes is very limited with five of the seven belonging to the Nosematida. Through multi-platform sequencing, we have effectively expanded upon existing genomes of Microsporidia sp. MB and the broader Enterocytozoonida with one of the first telomere-to-telomere, complete genome assembly in the clade. This novel complete assembly of MB SOUVK7 expands and diversifies current genomic datasets into the Mrazekiidae, whilst providing further insight into unique genetic characteristics of MB. Although the Microsporidia phylum exhibits a wide range of genome sizes (1.52-51.35 Mb, *Alternosema astaquatica* and *E. aedis* respectively), the 9.16 Mb genome of MB SOUVK7 represents a notable increase in genome size within the Enterocytozoonidae family, which have previously reported sizes of 6 Mb or lower (**Fig 3**). This includes a near 2-fold size increase compared to the 5.9 Mb assembly of Microsporidia sp. MB AHL03 whilst reducing contig number from 2335 to 13 [9,19]. Despite this increase in size, the predicted number of ORFs was similar to that of closely related genomes, which suggests that much of the size increase may be attributed to the correct assembly of intergenic repeat regions associated with longer reads. A prediction of 28.2% interspersed repeats in MB SOUVK7, compared to reported predictions of 0.57% or 7.32% mean in the Enterocytozoonidae (St Dev: 4.45%) supports this [9,19,20]. The MB SOUVK7 genome also contains a shift in proportion of transposable elements (TEs) away from DNA transposons (2.03% and 5.83%) and towards LINEs (24.40% and 0.34%, % assembly for MB SOUVK7 and mean Enterocytozoonidae). Although longer and more abundant LINE TEs are revealed by complete assembly, it is also possible that the symbiotic nature and principally vertical transmission of MB may have some influence on the evolution and size of the MB genome [5,21,22]. Vertical transmission and asexual reproduction often lead to strong genetic bottlenecks, resulting in expansions of LTR and LINE transposable elements within a population. This could explain genome size expansion in several vertically-transmitted Microsporidia [20,23]. Alternatively, the expansion in transposable elements could be the result of relaxed selection and the inability to eliminate mildly-deleterious transposable element insertions.

Flanking sequences of MB chromosomes provide further evidence of telomeric evolution across the Microsporidia with the TTAG motif observed in MB SOUVK7 representing the most reduced repeat in the phylum. Two guanine deletion events within the telomeric regions are evident across the phylum, firstly at the node shared with *Hamiltosporidium* species (TTAGGG to TTAGG) and secondly at the node shared with the Nosematida family (TTAGG to TTAG). The recently released genome of *Glugoides intestinalis* also shows a 4-mer TTAG repeat motif, further supporting telomeric motif reduction within the Enterocytozoonida [24,26]. No telomeric motifs are reported for *A. locustae,* despite its chromosome level assembly, which could otherwise better localise the taxonomic position for the first reduction event [27]. Telomeric simple repeats in MB SOUVK7 are also flanked by short stretches of LINE repeats of up to 10kb in the majority of chromosomes. In *Encephalitozoon* assemblies, telomeric CpG methylation is more frequent than in the rest of the genome, but the reverse is seen in MB SOUVK7 [14]. Synteny analysis shows no evidence of subtelomeric regions, which are characterised by mosaic collections of interchangeable segmental duplications (EXT repeats), despite their detection in all *Encephalitozoon* species [28]. Although subtelomeres appear to be lacking in MB, the presence of highly repetitive LINEs adjacent to the telomeric regions may echo subtelomeric characteristics without the broader homology between chromosomes.

In addition to telomeres, all 13 chromosomes exhibit distinct regions of differential genomic architecture, with several characteristics correlating with centromeric regions in fungi and broader eukaryotes. These regions are at least 10kb from chromosome ends and are gene sparse and GC rich whilst possessing dense clusters of CpG methylation and specific retroelements. These regions were identified by localisation of non-R4 LINE and LTR retroelements, which only represent approximately 3.7% and 3.4% of total retroelement predictions. Despite lacking variety these elements still represent 18.7% of bases associated with repeats. These repeat clusters have knock-on effects on local GC content and gene density. Ty3 LTRs are found almost exclusively in these regions and have been identified within centromeres of plants and fungi [29,30]. Microsporidian Ty3’s have however been found to have diversified dramatically within the phylum with the typical two ORFs of pol and gag being amalgamated into a single polyprotein [31]. The nearby presence of large rRNAs to these distinct regions in MB SOUVK7 may mimic the roles of long non-coding RNAs (lncRNAs), similar to mammalian and fungal centromeres [32]. Changes in local GC content can also define centromeres, however in fungal centromeres and similar regions in *A. locustae,* a local reduction in GC content is seen contrary to the increase observed in MB [25,32,33]. In MB SOUVK7 the GC content of these regions is actually higher than or equal to the GC content of other microsporidian genomes, with some MB regions having up to 49.5% GC. It must be noted however, that total mean GC content of both *A. locustae* and higher fungi are considerably greater than that of MB (41.6% and 40-65%, compared to 31.13% respectively), which may limit the biological feasibility of further GC reduction in MB [27,29].

Finally, ONT analysis of these regions consistently demonstrate elevated CpG methylation compared to the rest of the chromosome. The prediction of many methylation-associated proteins in MB supports the functionality of the CpG methylation pathways. Highly localised methylation is observed in centromeric regions of fungi, plants and animals, supressing gene expression and assisting CENP-A and H3 nucleosomes with centromeric chromatin formation [33]. Methylation of microsporidian chromosomes has only been examined in the Encephalitozoonidae, showing elevation in the telomeric and sub-telomeric regions, however this may not be comparable due to their lack of regional centromeres [14]. Within the broader fungal kingdom, elevated CpG methylation is colocalised with the peripheries of sites of CenH3 (CENP-A ortholog) binding in *Neospora crassa* and *Vesticillium dahliae*, whilst also correlating with low local gene density and elevated repeat density [29,34]. Although more data is required to confirm the final positions of centromeres in any microsporidian species, we note that the regions of interest highlighted by this study exhibit many of the characteristics of centromeres observed in broader eukaryotes, and hypothesize some microsporidia species may possess classical centromeres, instead of the point centromeres previously reported in the Encephalitozoonidae [14].

Predictions of MB tetraploidy agrees with the prior association of high ploidy species with vertical transmission and insect infection [23,35]. In a recent ploidy screen of 16 species, tetraploidy was identified in 6 species from a broad range of families (Neopereziida, Glugeida, Nosematida and Enterocytozoonida), whilst still possessing other diploid members in their respective families. Although not mentioned in the article, all tetraploid species are known to infect a variety of freshwater arthropod hosts, whilst 8 of the remaining diploid species infect hosts of other phyla. This association is consolidated by the tetraploid characterisation of *V. necatrix*, and several novel uncharacterised assemblies from arthropod hosts exhibiting polyploidy (14 of 19 assemblies)[25,36]. All ploidy analysis demonstrates a clear favoured kmer distribution of AAAB in MB, which also correlates with previously analysed microsporidia, supporting the broader trend of the phylum. Furthermore, predicted proportions of these kmer patterns in MB were found to be within the lower end of predicted values across the Microsporidia, suggesting relatively low levels of heterozygosity. This may stem from genetic bottlenecks associated with the primarily vertical transmission of MB, forcing populations within individual hosts to become relatively homogenous.

The proteins of microsporidian polar tube and spore wall have been shown to play important roles in both colonising and manipulating the host cells [1,17,37]. Although some protein families are well conserved throughout the phylum, other SWPs and PTPs have been found to be unique to specific genera and families. Given the loss and reduction of metabolic genes within the Enterocytozoonida, a potential minimalization of infection machinery was examined within the clade [18]. Our analysis found loss of PTP1, PTP4, PTP6, EnP1 and SWP9 in MB SOUVK7. The family-wide lack of PTP1 and PTP4 orthologs are surprising given their integral roles in polar tube adhesion to the host cell and previous electron micrographs demonstrating typical polar tubes within MB spores [22,37,38]. Notably, PTP1 and PTP4 both associate closely to the paralogous proteins of PTP2 and PTP5, respectively, and likely stem from recent gene duplication events [38–40]. As the presence of PTP2 proteins and polar tube functionality has been experimentally confirmed in the Enterocytozoonidae, this may suggest reduction events of PTP1 and PTP4 within this clade whilst the remaining paralogs have retained organelle functionality Finally the loss of PTP6 in the Mraezekiidae marks another reduction in polar tube components, however the primarily vertical lifecycle of MB may limit the necessity for a diverse variety of host invasion proteins [21,22,41]. SWP9 was also found to be lacking in all species of the Enterocytozoonida, mirroring the distribution of PTP1 and PTP4. SWP9 has been found to interact directly with the polar tube and PTP1 specifically, potentially as a polar tube anchoring component [42]. Given the proposed loss of PTP1 through gene reduction, the loss of SWP9 is feasible due to functional redundancy.

Whilst MB SOUVK7 exhibits many examples of gene redundancies, SWP12 was found to be a potential instance of gene diversification with two in-paralogs being observed across the Enterocytozoonida clade. Similar to SWP9, SWP12 contains a Bin/Amphiphysin/Rvs (BAR) domain, enabling host cell adhesion through GAG-binding [42,43]. SWP12 may also assist in parasitism beyond adhesion, playing roles in lipoprotein binding, nutrient sequestering and vacuole formation during merogony and sporogony [44]. The functional overlap of SWP12 may have enabled SWP9 and EnP1 gene reduction without impacting parasitic pathways or MB retention in the host. As all experimental data for SWP12 is currently in *N. bombycis*, further characterisation of these predicted paralogs within the Enterocytozoonida may clarify their function further elucidate any deviations in SWP12 function.

With the recent description of MB and its potential as an agent for malaria control, the Enterocytozoonida are receiving renewed attention [5,21]. As one of the first complete genomes within the family, the release of the MB SOUVK7 assembly expands our understanding of genomic architecture within the Enterocytozoonida and may act as a complete reference for population studies. MB has been identified in multiple African countries and several *Anopheles* species, with phenotypic differences already being noted between populations [4,6,7,21,22]. Further sequencing and assembly of complete MB genomes, whether *de novo* or assembled against MB SOUVK7, will better highlight genomic variance between populations and identify genes associated with host tropism, regional diversity and *Plasmodium* inhibition. Other mosquito-infecting microsporidia, including *V. culicus,* have previously been considered as agents for malaria control, however high levels of host pathogenicity during horizontal transmission have limited their potential for environmental dispersion and retention [45]. As one of the only Enterocytozoonida species to exhibit a primarily vertical lifecycle, genetic comparisons of MB to species in the wider family may also highlight genes and pathways that enable this lifestyle. [45] The near-symbiotic life cycle of MB with anopheline mosquitoes is a crucial factor in its suitability as a control agent and so better understanding its mechanisms for limiting host parasitism may further help in developing MB dispersal protocols.

## Materials and Methods

### Mosquito rearing and material isolation

Microsporidia sp. MB positive material was isolated from a previously lab-adapted *Anopheles coluzzii* colony, originating from Burkina Faso. Rearing and sampling for this study was performed at the MRC Centre for Virus Research, University of Glasgow UK. Mosquitoes were maintained at 28 °C at a constant 80 % relative humidity with 12-hour day/night light cycles. Early instar larvae were fed on liver powder (Now Foods) and progressed to TabiMin tablets (Tetra) from second instar. Adults were reared in cages of 500-2000 individuals and provided 5% sucrose solution *ad libitum*. Four days post-eclosure, adult females were fed with human blood (Scottish National Blood Transfusion Service, Glasgow) administered through a membrane feeder system (Hemotek ltd) and males and non-blood fed females were removed. Moist filter paper cones were placed into darkened cages 96 hours post-bloodmeal to promote immediate egg laying behaviour.

### DNA isolation and sample verification

PacBio sequencing was performed on dissected ovaries from gravid females, 96 hours post bloodmeal. Females were anaesthetised through cold exposure and gravid ovaries were dissected individually, with PBS being changed between dissections. Ovaries were pooled and transferred to –20 °C for storage. DNA extraction was performed on 100 ovary pairs, using the Gentra Puregene Tissue kit (4 g; QIAGEN) following the provided ‘DNA Purification from Tissue’ protocol. The final high molecular weight (HMW) DNA sample was eluted into 50 μl of molecular-grade water and stored at 4 °C until submission to Edinburgh Genomics, University of Edinburgh.

Nanopore sequencing provided additional long-read sequence and CpG methylation data for post-assembly analysis. As with the PacBio sample, 100 gravid ovary pairs were dissected, pooled in PBS and stored at –20 °C prior to extraction. Sample DNA was extracted with the Blood & Cell Culture DNA Kit (QIAGEN) following the provided Genomic DNA Preparation methods for 20/G columns. Extracted HMW samples were eluted into 50 μl of molecular-grade water and stored at 4 °C until submission to Edinburgh Genomics, University of Edinburgh.

Illumina sequencing for ploidy analysis was performed on sterilised, early-stage eggs to maximise the Microsporidia MB-host DNA ratio. To ensure the minimal age of the embryos, egg cones were swapped within gravid female cages hourly for six consecutive hours. Collected eggs were covered and allowed to melanise at room temperature for one hour, before being inactivated at –20 °C for a minimum of one hour. All inactivated egg cones were pooled and stored at –20 °C until sufficient material was collected. Once approximately 300-500 eggs were collected, all egg cones were thawed, and eggs were washed onto a vacuum membrane filter. The eggs were sterilised for five minutes by submersion in 1% hypochlorite solution (1:10 10 % hypochlorite in water), drained through vacuum filtration and repeatedly washed in sterile PBS for an additional 5 minutes. Eggs were transferred to a 1.5 ml centrifuge tube and DNA was extracted through a modified Phenol:Chloroform extraction method. Briefly, all liquid was removed from the egg suspension and 500 μl of phenol was added to the centrifuge tube. All eggs were ruptured through manual grinding with a micropestle, before adding 500μl of chloroform and mixing through inversion. The extraction mix was centrifuged (13,000 x g for 10 minutes at 4 °C), the aqueous layer was collected and the DNA was precipitated with an equal volume of precipitation solution (40 µl 3 M sodium acetate in 800 µl absolute ethanol) for 60 minutes at –20 °C. Precipitated DNA was collected through centrifugation (13,000 g for 10 minutes at 4 °C), washed in 70 % ethanol, eluted into 100 μl molecular-grade water and stored at 4 °C until submission to Glasgow Polyomics (now MVLS Shared Research Facilities), University of Glasgow UK.

### Genome sequencing and assembly

A single HMW DNA sample was submitted for PacBio sequencing, with sample quality and average DNA fragment size being assessed through TapeStation (Agilent Technologies ltd), Femto Pulse (Agilent Technologies ltd) and dsDNA QuBit (ThermoFisher Scientific Inc) systems. The sample library was prepared with 10.95 μg of HMW gDNA and sequenced with a Revio SMRTbell prep kit on a Revio SMRT 25 M ZMW cell using HiFi mode (PacBio). Initial read quality and distributions were verified with FastQC (https://www.bioinformatics.babraham.ac.uk/projects/fastqc). Host contamination was removed by mapping the Pacbio HiFi reads to an *An. gambiae* reference genome (NCBI accession: GCA_943734735.2) with minimap2 v2.17 [46]. Decontaminated reads were initially assembled with Hifiasm v0.19.8, following default parameters [47]. The taxonomy profiles and GC content of the assembled contigs were analysed with SprayNPray and a custom database of microsporidian, bacterial and non-microsporidian eukaryotic genomes [48]. Sequencing depth of assembled contigs was calculated with Samtools ‘depth’ command [49]. After the prediction of potential tetraploidy of MB, a final genome was reassembled using Hifiasm with ploidy set to 4 (--nhap option).

A single Illumina sample was submitted, NGS library prepared with 9.1 μg of gDNA using the NEBNext Ultra II FS DNA Library Prep Kit for Illumina (New England Biolabs) and sequenced on a NextSeq2000 with 2*100 bp paired-end reads (Illumina Inc). The provided adapter sequences were trimmed with Trimmomatic v0.39 [50] and host reads were removed using the *An. gambiae* genome with Bowtie2 v2.4.2 [51].

Nanopore sequencing was prepared with 1.5 μg of HMW gDNA using a Ligation Sequencing Kit V14 and sequenced with a PromethION system (Oxford Nanopore Technologies plc, UK). Data was provided by Edinburgh Genomics as .fastq and .bam files for the raw reads and methylation base calls, respectively. The methylation BAM file was aligned to the final MB SOUVK7 assembly with Dorado 0.5.2 aligner, reviewed with Modkit 0.4.4 summary function with ‘no filtering’ and the final bedMethyl file of base modification calls was generated with Modkit pileup using the “traditional” preset (https://github.com/nanoporetech). Methylation density was calculated across all chromosomes as the frequency of CpG methylation over total bases within 3,000 bp windows.

### Gene prediction and genome annotation

Genome masking was performed with RepeatModeler (https://github.com/Dfam-consortium/RepeatModeler) and RepeatMasker (https://github.com/Dfam-consortium/RepeatMasker) for repeat family identification and genome masking, respectively. To ensure prediction of broader microsporidian repeat families, modelling was performed on the chromosome-level genomes of *A. locustae* CLX, *E. cuniculi* GB-M1, *H. tvaerminnensis* FI-OER-3-3, *V. necatrix* (NCBI accession: GCA_007674295.1, GCF_000091225.2, GCA_022605425.2 and GCF_036630325.1, respectively) and the final assembly of Microsporidia sp. MB SOUVK7. Gene prediction and genome annotation was performed with Funannotate (https://github.com/nextgenusfs/funannotate), supplementing prediction with independently run Augustus v3.5.0 [52] and GeneMark-ES [53] files. Augustus gene prediction was run specifying species as ‘encephalitozoon_cuniculi_GB’ and ‘intronless’ for gene model parameters and GeneMark-ES 2.5 was run following eukaryotic prediction (‘-euk’). Funannotate ‘predict’ function was run using the ‘microsporidia_odb10’ BUSCO database, whilst supplementing the previous prediction files and removing GlimmerHMM functionality from the pipeline, due to excessive gene fragmentation. InterProScan analysis was run locally as part of the Funannotate ‘iprscan’ command, with resulting files being provided as part of ‘annotate’ command along with the ‘microsporidia_odb10’ BUSCO database.

Telomeric regions were identified from the results of RepeatMasker, verifying the presence of (TTAG)_n_ repeat regions at the ends of each chromosome. Repeat content along MB chromosomes were incorporated and visualized in Circos as custom BED files, using the shinyCircos-V2 R package [54]. Final genome integrity was assessed with BUSCO v5.4.5 against the microsporidia_odb12 dataset [55].

As ribosomal RNA (rRNA) prediction is currently not implemented into the Funannotate pipeline, rRNA prediction was performed with Barrnap 0.9 (https://github.com/tseemann/barrnap), specifying bacteria for the kingdom parameter to account for the reduced size of microsporidian rRNAs. The resulting GFF file was integrated into the primary Funannotate output GFF with AGAT ‘manage_IDs’ tool (https://github.com/NBISweden/AGAT).

### Genome ploidy, gene orthology and GO analysis

For comparison of MB SOUVK7 assembly quality to other published genomes, statistics of NCBI reference genomes were collected with a focus on assemblies of higher levels (“Complete”/“Chromosome”) and closely related species (S3 Table).

To initially assess the haploid assembly for potential polyploidy, chromosomal synteny was examined with MCScanX [56]. Host-decontaminated reads of both Illumina and PacBio samples that mapped to the final MB assembly were also used to analyse ploidy. For reference-free kmer analysis, GenomeScope 2.0 was run using 21 bp kmer length, and visualised with Smudgeplot, following default parameters [57]. Additional validation of predicted ploidy was provided with ploidyNGS V3.1.3 [11].

Predicted genes of MB SOUVK7 were further investigated to assess gene orthology, encoded-protein function and localisation, and genetic domains. Comparative orthology between MB SOUVK7 and all other NCBI available proteomes of Microsporidia (46 proteomes, S3 Table) was assessed using Orthofinder v2.5.5 [58], including *M. daphniae* and *Amphiamblys sp.* acting as an outgroup as members of the Metchnikovellids [59]. Examination of all to all BLASTp and tBLASTn search results confirmed the lack of orthologs in any species. BLAST predictions were only included into results if all BLAST-predicted orthologs were identified with an e-value > 1e^-5^ and existed in an orthogroup containing all undetected orthologs. Protein function was predicted with stand-alone DeepLoc2.0 [60] using the ‘accurate’ methodology. Additional genetic domains were predicted with standalone versions of TargetP-2.0 [61] and DeepTMHMM [62], to predict signal peptides and transmembrane domains, respectively.

To examine the presence or absence of polar tube and spore proteins in MB SOUVK7 and broader Microsporidia, the orthogroup results of the above OrthoFinder analysis were searched against a pre-determined list of confirmed PTP and SWP proteins (S5 Table). Orthogroups with corresponding orthologs were compiled, with entries being concatenated when multiple search orthologs for the same protein family was provided and multiple orthogroups were found. Any consistent lack of orthologs across both MB genomes or Enterocytozoonida was verified through tBLASTn searches of all identified orthologs to the relative genomes.

Genes associated with methylation, telomeric and centromeric maintenance were previously reported in the Encephalitozoonidae [14] and so potential orthologs in MB were identified to hits in *Encephalitozoon cuniculi* using BLASTP with an e-value threshold of 1E^-5^. Given the reduced nature of the Encephalitozoonidae genomes, genes associated to gene ontology (GO) terms for telomere maintenance (GO:0000723), heterochromatin formation (GO:0031507), centromere complex assembly (GO:0034508) and methylation (GO:0032259) for *Allomyces macrogynus, Mucor lusitanicus, Rozella allomycis, Saccharomyces cerevisiae* and *Schizosaccharomyces pombe* were supplemented into the existing OrthoFinder dataset (NCBI accession:: GCA_000151295.1, GCA_001638945.1, GCA_000442015.1, GCA_000146045.2 and GCA_000002945.3, respectively). Orthogroups including any of the supplemented GO genes were extracted from the total orthogroup data frame, and potential microsporidian orthologs were examined.

## Availability of Data and Materials

The datasets supporting the conclusions of this article are included within the article (and its additional files) and available in the NCBI BioProject repository, [PRJNA1200702 & PRJNA1211496]. The final assembly sequence of Microsporidia sp. MB SOUVK7 and associated feature files were submitted to NCBI WGS database under the accession SUB14942438. All raw Illumina, PacBio and ONT Nanopore read data generated in this study were submitted to the NCBI Sequence Read Archive (SRA; https://www.ncbi.nlm.nih.gov/sra) under accession numbers: SRR32024117, SRR32024118 and SRR35190449, respectively

## Acknowledgements

We thank the teams at MVLS Shared Research Facilities and Edinburgh Genomics for assistance and feedback for optimising sequencing samples and extraction methods for their respective platforms. The authors would also like to thank Lilian Ang’ang’o and Jeremy Herren (*icipe,* Kenya) for sharing and discussing their MB assemblies prior to publication, along with ongoing communications between our groups.

## Supporting Information

**S1_Fig.jpg:**

**S1 Fig**: Overview of reads and haploid assembly of PacBio sequencing dataset, before and after host read decontamination. A. FastQC overview of Raw PacBio read distributions of read count against read GC content. Two distinct peaks are visible of approximately 300,000 and 420,000 reads with GC content of 31% and 45%, respectively. B. FastQC overview of PacBio reads after removal of reads aligned to An. gambiae, exhibiting only a single peak of approximately 300,000 reads and 31% GC content. C. SprayNPray output figure of contigs assembled from host decontaminated reads following haploid assembly. The predicted phylogenetic origins of contigs are illustrated through point colour, with uncharacterised and bacterial contigs being clustered with low read depth and high GC content. Contigs for further syntenic alignment were isolated for the elongated cluster of 55 contigs with a GC content of ∼31% and >40x sequencing depth.

**S2_Table.xlsx:**

**S2 Table**: Tabulated output of RepeatMasker run against Microsporidia sp. MB SOUVK7

**S3_Table.xlsx:**

**S3 Table**: Microsporidia assemblies included in the comparison on Microsporidia sp. MB SOUVK7 against the rest of the phylum. M. daphniae and Amphiamblus sp. was included as outgroups as a basal species of the Microsporidia and Cryptomycota, respectively.

**S4_Table.xlsx:**

**S4 Table**: List of signal peptide and transmembrane domain prediction of genes in Microsporidia sp. MB, including consensus gene counts between predictive tools

**S5_Table.xlsx:**

**S5 Table**: List of predicted orthologs to polar tube and spore wall proteins across species of the Microsporidia, as illustrated in ***Fig 4***

## References

1. Han B, Takvorian PM, Weiss LM. Invasion of Host Cells by Microsporidia. Front Microbiol. 2020;11. doi:10.3389/fmicb.2020.00172

2. Stentiford GD, Becnel JJ, Weiss LM, Keeling PJ, Didier ES, Williams BAP, et al. Microsporidia – Emergent Pathogens in the Global Food Chain. Trends Parasitol. 2016;32: 336–348. doi:10.1016/j.pt.2015.12.004

3. Ruan Y, Xu X, He Q, Li L, Guo J, Bao J, et al. The largest meta-analysis on the global prevalence of microsporidia in mammals, avian and water provides insights into the epidemic features of these ubiquitous pathogens. Parasit Vectors. 2021;14: 186. doi:10.1186/s13071-021-04700-x

4. Akorli J, Akorli EA, Tetteh SNA, Amlalo GK, Opoku M, Pwalia R, et al. Microsporidia MB is found predominantly associated with Anopheles gambiae s.s and Anopheles coluzzii in Ghana. Sci Rep. 2021;11: 18658. doi:10.1038/s41598-021-98268-2

5. Herren JK, Mbaisi L, Mararo E, Makhulu EE, Mobegi VA, Butungi H, et al. A microsporidian impairs Plasmodium falciparum transmission in Anopheles arabiensis mosquitoes. Nat Commun. 2020;11: 2187. doi:10.1038/s41467-020-16121-y

6. Tchigossou G, Lontsi-Demano M, Tossou E, Sovegnon P-M, Akoton R, Adanzounon D, et al. Seasonal variation of Microsporidia MB infection in Anopheles gambiae and Anopheles coluzzii in two different geographical localities in Benin. Malar J. 2025;24: 95. doi:10.1186/s12936-025-05247-3

7. Moustapha LM, Sadou IM, Arzika II, Maman LI, Gomgnimbou MK, Konkobo M, et al. First identification of Microsporidia MB in Anopheles coluzzii from Zinder City, Niger. Parasit Vectors. 2024;17: 39. doi:10.1186/s13071-023-06059-7

8. Nakjang S, Williams TA, Heinz E, Watson AK, Foster PG, Sendra KM, et al. Reduction and expansion in microsporidian genome evolution: new insights from comparative genomics. Genome Biol Evol. 2013;5: 2285–2303. doi:10.1093/gbe/evt184

9. Ang’ang’o LM, Waweru JW, Makhulu EE, Wairimu A, Otieno FG, Onchuru T, et al. Draft genome of Microsporidia sp. MB—a malaria-blocking microsporidian symbiont of the Anopheles arabiensis. Microbiol Resour Announc. 2024;13: e00903–23. doi:10.1128/MRA.00903-23

10. Bojko J, Reinke AW, Stentiford GD, Williams B, Rogers MSJ, Bass D. Microsporidia: a new taxonomic, evolutionary, and ecological synthesis. Trends Parasitol. 2022;38: 642–659. doi:10.1016/j.pt.2022.05.007

11. Augusto Corrêa dos Santos R, Goldman GH, Riaño-Pachón DM. ploidyNGS: visually exploring ploidy with Next Generation Sequencing data. Bioinformatics. 2017;33: 2575–2576. doi:10.1093/bioinformatics/btx204

12. Badie S, Escandell JM, Bouwman P, Carlos AR, Thanasoula M, Gallardo MM, et al. BRCA2 Acts as RAD51 Loader to Facilitate Telomere Replication and Capping. Nat Struct Mol Biol. 2010;17: 1461–1469. doi:10.1038/nsmb.1943

13. Verma P, Dilley RL, Zhang T, Gyparaki MT, Li Y, Greenberg RA. RAD52 and SLX4 act nonepistatically to ensure telomere stability during alternative telomere lengthening. Genes Dev. 2019;33: 221–235. doi:10.1101/gad.319723.118

14. Mascarenhas dos Santos AC, Julian AT, Liang P, Juárez O, Pombert J-F. Telomere-to-Telomere genome assemblies of human-infecting Encephalitozoon species. BMC Genomics. 2023;24: 237. doi:10.1186/s12864-023-09331-3

15. Kuscu C, Zaratiegui M, Kim HS, Wah DA, Martienssen RA, Schalch T, et al. CRL4-like Clr4 complex in Schizosaccharomyces pombe depends on an exposed surface of Dos1 for heterochromatin silencing. Proc Natl Acad Sci U S A. 2014;111: 1795–1800. doi:10.1073/pnas.1313096111

16. De Rop V, Padeganeh A, Maddox PS. CENP-A: the key player behind centromere identity, propagation, and kinetochore assembly. Chromosoma. 2012;121: 527–538. doi:10.1007/s00412-012-0386-5

17. Weiss LM, Delbac F, Russell Hayman J, Pan G, Dang X, Zhou Z. The Microsporidian Polar Tube and Spore Wall. Microsporidia. John Wiley & Sons, Ltd; 2014. pp. 261–306. doi:10.1002/9781118395264.ch10

18. Wiredu Boakye D, Jaroenlak P, Prachumwat A, Williams TA, Bateman KS, Itsathitphaisarn O, et al. Decay of the glycolytic pathway and adaptation to intranuclear parasitism within Enterocytozoonidae microsporidia. Environ Microbiol. 2017;19: 2077–2089. doi:10.1111/1462-2920.13734

19. Ang’ang’o LM, Herren JK, Tastan Bishop Ö. Bioinformatics analysis of the Microsporidia sp. MB genome: a malaria transmission-blocking symbiont of the Anopheles arabiensis mosquito. BMC Genomics. 2024;25: 1132. doi:10.1186/s12864-024-11046-y

20. de Albuquerque NRM, Ebert D, Haag KL. Transposable element abundance correlates with mode of transmission in microsporidian parasites. Mob DNA. 2020;11: 19. doi:10.1186/s13100-020-00218-8

21. Onchuru TO, Makhulu EE, Ronnie PC, Mandere S, Otieno FG, Gichuhi J, et al. The Plasmodium transmission-blocking symbiont, Microsporidia MB, is vertically transmitted through Anopheles arabiensis germline stem cells. PLOS Pathog. 2024;20: e1012340. doi:10.1371/journal.ppat.1012340

22. Parry ERS, Pevsner R, Poulton BC, Purusothaman D-K, Adam AI, Issiaka S, et al. Imaging the lifecycle of Microsporidia sp. MB in Anopheles coluzzii from western Burkina Faso reveals octosporogony. mSphere. 2025;0: e00851–24. doi:10.1128/msphere.00851-24

23. Haag KL, Pombert J-F, Sun Y, de Albuquerque NRM, Batliner B, Fields P, et al. Microsporidia with Vertical Transmission Were Likely Shaped by Nonadaptive Processes. Genome Biol Evol. 2019;12: 3599–3614. doi:10.1093/gbe/evz270

24. Angst P, Pombert J-F, Ebert D, Fields PD. Near chromosome–level genome assembly of the microsporidium Hamiltosporidium tvaerminnensis. G3 GenesGenomesGenetics. 2023;13: jkad185. doi:10.1093/g3journal/jkad185

25. Svedberg D, Winiger RR, Berg A, Sharma H, Tellgren-Roth C, Debrunner-Vossbrinck BA, et al. Functional annotation of a divergent genome using sequence and structure-based similarity. BMC Genomics. 2024;25: 6. doi:10.1186/s12864-023-09924-y

26. Angst P, Ebert D, Fields PD. Variation in genome architecture and epigenetic modification across the microsporidia phylogeny. Genome Biol Evol. 2025; evaf166. doi:10.1093/gbe/evaf166

27. Chen L, Gao X, Li R, Zhang L, Huang R, Wang L, et al. Complete genome of a unicellular parasite (Antonospora locustae) and transcriptional interactions with its host locust. Microb Genomics. 2020;6: e000421. doi:10.1099/mgen.0.000421

28. Dia N, Lavie L, Faye N, Méténier G, Yeramian E, Duroure C, et al. Subtelomere organization in the genome of the microsporidian Encephalitozoon cuniculi: patterns of repeated sequences and physicochemical signatures. BMC Genomics. 2016;17: 34. doi:10.1186/s12864-015-1920-7

29. Seidl MF, Kramer HM, Cook DE, Fiorin GL, van den Berg GCM, Faino L, et al. Repetitive Elements Contribute to the Diversity and Evolution of Centromeres in the Fungal Genus Verticillium. mBio. 2020;11: 10.1128/mbio.01714-20. doi:10.1128/mbio.01714-20

30. Heuberger M, Koo D-H, Ahmed HI, Tiwari VK, Abrouk M, Poland J, et al. Evolution of Einkorn wheat centromeres is driven by the mutualistic interplay of two LTR retrotransposons. Mob DNA. 2024;15: 16. doi:10.1186/s13100-024-00326-9

31. Parisot N, Pelin A, Gasc C, Polonais V, Belkorchia A, Panek J, et al. Microsporidian Genomes Harbor a Diverse Array of Transposable Elements that Demonstrate an Ancestry of Horizontal Exchange with Metazoans. Genome Biol Evol. 2014;6: 2289–2300. doi:10.1093/gbe/evu178

32. Yadav V, Sreekumar L, Guin K, Sanyal K. Five pillars of centromeric chromatin in fungal pathogens. PLoS Pathog. 2018;14: e1007150. doi:10.1371/journal.ppat.1007150

33. Smith KM, Galazka JM, Phatale PA, Connolly LR, Freitag M. Centromeres of filamentous fungi. Chromosome Res Int J Mol Supramol Evol Asp Chromosome Biol. 2012;20: 635–656. doi:10.1007/s10577-012-9290-3

34. Smith KM, Phatale, Pallavi A., Sullivan, Christopher M., Pomraning, Kyle R., and Freitag M. Heterochromatin Is Required for Normal Distribution of Neurospora crassa CenH3. Mol Cell Biol. 2011;31: 2528–2542. doi:10.1128/MCB.01285-10

35. Khalaf A, Lawniczak MKN, Blaxter ML, Jaron KS. Polyploidy is widespread in Microsporidia. Microbiol Spectr. 2024;12: e03669–23. doi:10.1128/spectrum.03669-23

36. Khalaf A, Zhou C, Weber CC, Vancaester E, Sims Y, Makunin A, et al. Forty New Genomes Shed Light on Sexual Reproduction and the Origin of Tetraploidy in Microsporidia. bioRxiv; 2025. p. 2025.05.12.652816. doi:10.1101/2025.05.12.652816

37. Han B, Ma Y, Tu V, Tomita T, Mayoral J, Williams T, et al. Microsporidia Interact with Host Cell Mitochondria via Voltage-Dependent Anion Channels Using Sporoplasm Surface Protein 1. mBio. 2019;10: e01944–19. doi:10.1128/mBio.01944-19

38. Han B, Polonais V, Sugi T, Yakubu R, Takvorian PM, Cali A, et al. The role of microsporidian polar tube protein 4 (PTP4) in host cell infection. PLoS Pathog. 2017;13: e1006341. doi:10.1371/journal.ppat.1006341

39. Polonais V, Prensier G, Méténier G, Vivarès CP, Delbac F. Microsporidian polar tube proteins: Highly divergent but closely linked genes encode PTP1 and PTP2 in members of the evolutionarily distant *Antonospora* and *Encephalitozoon* groups. Fungal Genet Biol. 2005;42: 791–803. doi:10.1016/j.fgb.2005.05.005

40. Wang L, Lv Q, Meng X, Chen J, Wang Y, Pan G, et al. Identification and characterization polar tube protein 2 (PTP2) from *Enterocytozoon hepatopenaei* and its potential effect on shrimp microsporidian germination activity evaluation. Aquaculture. 2021;544: 737062. doi:10.1016/j.aquaculture.2021.737062

41. Lv Q, Wang L, Fan Y, Meng X, Liu K, Zhou B, et al. Identification and characterization a novel polar tube protein (NbPTP6) from the microsporidian Nosema bombycis. Parasit Vectors. 2020;13: 475. doi:10.1186/s13071-020-04348-z

42. Yang D, Pan L, Peng P, Dang X, Li C, Li T, et al. Interaction between SWP9 and Polar Tube Proteins of the Microsporidian Nosema bombycis and Function of SWP9 as a Scaffolding Protein Contribute to Polar Tube Tethering to the Spore Wall. Infect Immun. 2017;85: e00872–16. doi:10.1128/IAI.00872-16

43. Chen J, Li Z, Sheng X, Huang J, Sun Q, Huang Y, et al. Heterologous Expressed NbSWP12 from Microsporidia Nosema bombycis Can Bind with Phosphatidylinositol 3-Phosphate and Affect Vesicle Genesis. J Fungi. 2022;8: 764. doi:10.3390/jof8080764

44. Wang C, Yu B, Meng X, Xia D, Pei B, Tang X, et al. Microsporidian Nosema bombycis hijacks host vitellogenin and restructures ovariole cells for transovarial transmission. PLOS Pathog. 2023;19: e1011859. doi:10.1371/journal.ppat.1011859

45. Vyas-Patel N. The Suppression of Plasmodium berghei in Anopheles coluzzii infected later with Vavraia culicis. bioRxiv; 2023. p. 2023.02.05.527158. doi:10.1101/2023.02.05.527158

46. Li H. Minimap2: pairwise alignment for nucleotide sequences. Bioinformatics. 2018;34: 3094–3100. doi:10.1093/bioinformatics/bty191

47. Cheng H, Asri M, Lucas J, Koren S, Li H. Scalable telomere-to-telomere assembly for diploid and polyploid genomes with double graph. Nat Methods. 2024;21: 967–970. doi:10.1038/s41592-024-02269-8

48. Garber AI, Armbruster CR, Lee SE, Cooper VS, Bomberger JM, McAllister SM. SprayNPray: user-friendly taxonomic profiling of genome and metagenome contigs. bioRxiv; 2021. p. 2021.07.17.452725. doi:10.1101/2021.07.17.452725

49. Danecek P, Bonfield JK, Liddle J, Marshall J, Ohan V, Pollard MO, et al. Twelve years of SAMtools and BCFtools. GigaScience. 2021;10: giab008. doi:10.1093/gigascience/giab008

50. Bolger AM, Lohse M, Usadel B. Trimmomatic: a flexible trimmer for Illumina sequence data. Bioinformatics. 2014;30: 2114–2120. doi:10.1093/bioinformatics/btu170

51. Langmead B, Salzberg SL. Fast gapped-read alignment with Bowtie 2. Nat Methods. 2012;9: 357–359. doi:10.1038/nmeth.1923

52. Stanke M, Diekhans M, Baertsch R, Haussler D. Using native and syntenically mapped cDNA alignments to improve de novo gene finding. Bioinformatics. 2008;24: 637–644. doi:10.1093/bioinformatics/btn013

53. Lomsadze A, Ter-Hovhannisyan V, Chernoff YO, Borodovsky M. Gene identification in novel eukaryotic genomes by self-training algorithm. Nucleic Acids Res. 2005;33: 6494–6506. doi:10.1093/nar/gki937

54. Wang Y, Jia L, Tian G, Dong Y, Zhang X, Zhou Z, et al. shinyCircos-V2.0: Leveraging the creation of Circos plot with enhanced usability and advanced features. iMeta. 2023;2: e109. doi:10.1002/imt2.109

55. Manni M, Berkeley MR, Seppey M, Simão FA, Zdobnov EM. BUSCO Update: Novel and Streamlined Workflows along with Broader and Deeper Phylogenetic Coverage for Scoring of Eukaryotic, Prokaryotic, and Viral Genomes. Mol Biol Evol. 2021;38: 4647–4654. doi:10.1093/molbev/msab199

56. Wang Y, Tang H, DeBarry JD, Tan X, Li J, Wang X, et al. MCScanX: a toolkit for detection and evolutionary analysis of gene synteny and collinearity. Nucleic Acids Res. 2012;40: e49. doi:10.1093/nar/gkr1293

57. Ranallo-Benavidez TR, Jaron KS, Schatz MC. GenomeScope 2.0 and Smudgeplot for reference-free profiling of polyploid genomes. Nat Commun. 2020;11: 1432. doi:10.1038/s41467-020-14998-3

58. Emms DM, Kelly S. OrthoFinder: phylogenetic orthology inference for comparative genomics. Genome Biol. 2019;20: 238. doi:10.1186/s13059-019-1832-y

59. Park E, Poulin R. Revisiting the phylogeny of microsporidia. Int J Parasitol. 2021;51: 855–864. doi:10.1016/j.ijpara.2021.02.005

60. Thumuluri V, Almagro Armenteros JJ, Johansen AR, Nielsen H, Winther O. DeepLoc 2.0: multi-label subcellular localization prediction using protein language models. Nucleic Acids Res. 2022;50: W228–W234. doi:10.1093/nar/gkac278

61. Armenteros JJA, Salvatore M, Emanuelsson O, Winther O, Heijne G von, Elofsson A, et al. Detecting sequence signals in targeting peptides using deep learning. Life Sci Alliance. 2019;2. doi:10.26508/lsa.201900429

62. Hallgren J, Tsirigos KD, Pedersen MD, Armenteros JJA, Marcatili P, Nielsen H, et al. DeepTMHMM predicts alpha and beta transmembrane proteins using deep neural networks. bioRxiv; 2022. p. 2022.04.08.487609. doi:10.1101/2022.04.08.487609

