## Supplementary figures and images for "Telomere-to-telomere genome assembly of Microsporidia sp. MB, a microsporidian symbiont of *Anopheles coluzzii* isolated from Burkina Faso"

### Supplemental Figure 1

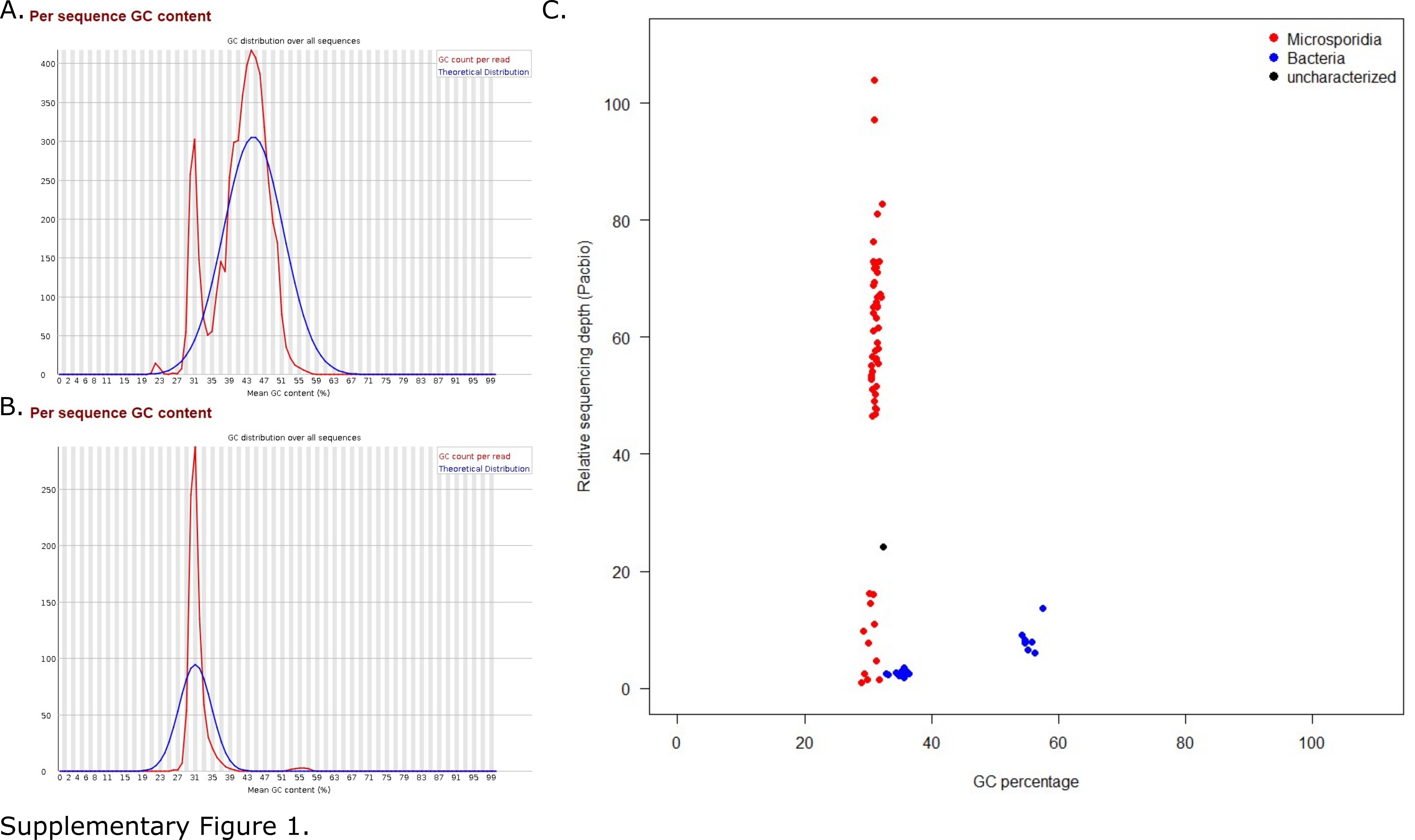
